# *dnmt1* function is required to maintain retinal stem cells within the ciliary marginal zone of the zebrafish eye

**DOI:** 10.1101/2020.01.29.925784

**Authors:** Krista M. Angileri, Jeffrey M. Gross

## Abstract

The ciliary marginal zone (CMZ) of the zebrafish retina contains a population of actively proliferating resident stem cells, which generate retinal neurons throughout life. The maintenance methyltransferase, *dnmt1*, is expressed within the CMZ. Loss of dnmt1 function results in gene misregulation and cell death in a variety of developmental contexts, however, its role in retinal stem cell (RSC) maintenance is currently unknown. Here, we demonstrate that zebrafish *dnmt1^s872^* mutants possess severe defects in RSC maintenance within the CMZ. Using a combination of immunohistochemistry, *in situ* hybridization, and a transgenic reporter assay, our results demonstrate a requirement for dnmt1 activity in the regulation of RSC proliferation, gene expression and in the repression of endogenous retroelements (REs). Ultimately, cell death is elevated in the *dnmt1*^-/-^ CMZ, but in a *p53*-independent manner. Using a transgenic reporter for RE transposition activity, we demonstrate increased transposition in the *dnmt1*^-/-^ CMZ. Taken together our data identify a critical role for dnmt1 function in RSC maintenance in the vertebrate eye.

## Introduction

The distal region of the vertebrate retina, termed the ciliary marginal zone (CMZ), contains a population of resident retinal stem cells (RSCs). The CMZ remains proliferative throughout the life of fish, but it proliferates to a more limited extent during the lifetime of amphibians and birds^1–6^. Whether an analogous structure exists in mammals is debated, but there are distinct, progenitor-like cells in the periphery of the retina that are active during embryogenesis^7–9^. Mammalian RSCs can also be isolated from the adult ciliary margin, cultured *in vitro*, and stimulated to produce retinal neurons^10–13^. However, this activity has not been demonstrated in the mature mammalian retinae *in vivo*.

Studies of the CMZ have primarily focused on zebrafish and *Xenopus* models to determine genetic pathways required for RSC identity^2,14–16^ and to characterize the epigenetic networks which regulate RSC function^17,18^. By comparison, the mechanisms mediating RSC maintenance *in vivo* remain unknown. In studies of RSCs, the zebrafish has been advantageous given that it possesses a highly active RSC population and is tractable for genetic and pharmacological manipulations, transgenesis and *in vivo* imaging^19,20^.

DNA methylation, a frequently studied epigenetic modification, is the process through which a methyl group is added to the fifth carbon of cytosine nucleotides and is commonly found at CpG dinucleotide sequences^21^. Members of the family of DNA methyltransferase (Dnmt) enzymes^22,23^ catalyze this epigenetic modification. Dnmt1 serves as a maintenance methyltransferase, copying the methylation pattern from parent to daughter strand during DNA replication and its function is required for cell cycle progression^24–26^. Loss of Dnmt1 function results in genomic hypomethylation^27–29^ and in developmental contexts and specific organ systems, this often compromises progenitor cell maintenance^24,27,30–33^ through numerous cellular mechanisms. These include: inducing cell cycle arrest^34,35^, retroelement activation^36–39^, inflammatory responses^33,37,40^, aberrant differentiation^28,31,41–44^ and/or *p53*-mediated apoptosis^34,35^.

Utilizing the *dnmt1^s872^* mutant zebrafish allele^30^, we establish an *in vivo* requirement for dnmt1 in RSCs. Through our analyses, we identify a decrease in overall RSC numbers, reduced RSC proliferation and aberrant gene expression patterns within the dnmt1-deficient CMZ. Additionally, we note increased retroelement expression and increased retrotransposition activity in *dnmt1*^-/-^ embryos. Remarkably, RSCs in *dnmt1*^-/-^ embryos are eliminated in a p53-independent manner, suggesting that dnmt1 represses alternative, non-apoptotic cell death pathways in RSCs. Taken together, these data highlight a novel function for dnmt1 in maintaining stem cell populations in the vertebrate retina.

## Results

### *dnmt1* mutants possess defects in the ciliary marginal zone

Previously, we identified a requirement for dnmt1 in maintaining lens epithelial cell viability using *dnmt1^s872^* mutant zebrafish^27^. During these previous studies, we also detected photoreceptor layer abnormalities, similar to those documented in *Dnmt1*^-/-^ conditional knockout mice^45,46^, and an apparent defect in the CMZ. With an interest in the role that dnmt1 plays in maintaining RSCs *in vivo*, here, we focused further on the CMZ phenotype. Using DAPI to label and count retinal nuclei, we confirmed a progressive degeneration of CMZ morphology beginning at 4 days post fertilization (dpf; Figure 1A-F) and a significant decline in retinal cell numbers through 5dpf (Figure 1G). The total number of cells present within central retina sections are equivalent between *dnmt1*^-/-^ and sibling larvae at 3dpf; however, numbers in *dnmt1*^-/-^ larvae diminish significantly between 4 and 5dpf (18.8% and 26.6% reduction respectively; *p* <0.0005; Figure 1G). Additionally, we compared the proportions of nuclei within the ganglion cell layer (GCL), inner nuclear layer (INL), outer nuclear layer (ONL), and CMZ between *dnmt1*^-/-^ larvae and siblings from 3-5dpf (Figure 1H). Interestingly, the proportions of cells in all three retinal laminae (GCL, INL, & ONL) remained equivalent over time in *dnmt1*^-/-^ larvae when compared to siblings, with only a slight increase in the ONL at 4dpf (Figure 1H and Supplementary Figure 1A-C; *p* <0.005). In contrast, the CMZ proportion decreased significantly from 3-5dpf suggesting that dnmt1 function in the retina is required within the CMZ to maintain the RSC population (Figure 1H and Supplementary Figure 1D; *p* <0.0005).

**Figure 1.**
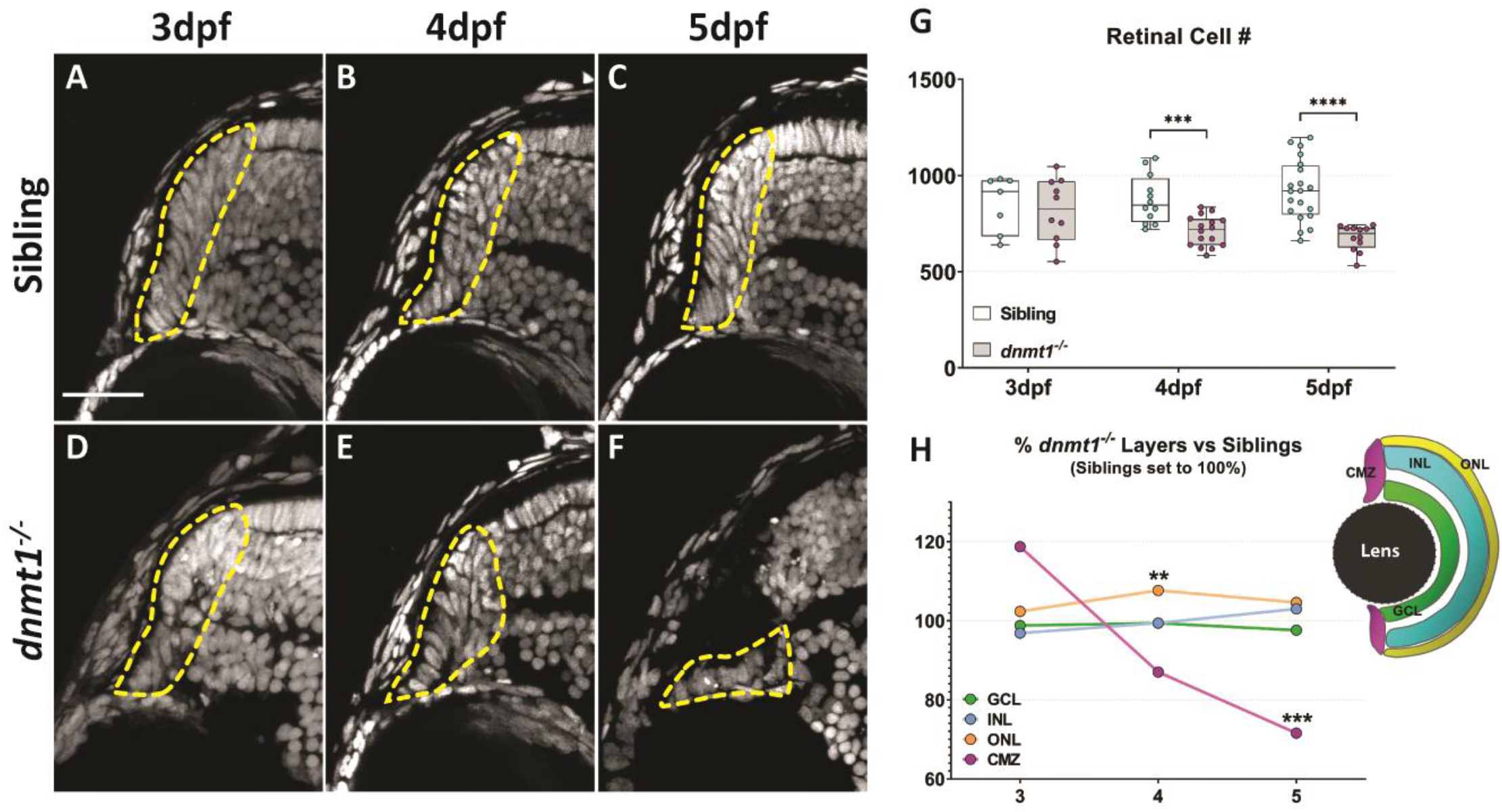
Disruption of dnmt1 function results in CMZ defects. A-F. DAPI staining of nuclei (gray) within the CMZ (white dotted lines delineate CMZ boundaries) of siblings (**A-C**) and *dnmt1*^-/-^ (**D-F**) larvae from 3-5dpf. **G.** Average number of all nuclei within the central retina of siblings and mutants. Each data point is the average of cell counts from three different 12 μm sections in one eye of a single larva. **H.** Proportional changes of *dnmt1*^-/-^ retinal domains (GCL, INL, ONL, & CMZ) relative to siblings (set to 100%). Colors correspond with retinal domains in diagram. Scale bars = 30 μm. \*\**p* <0.005, *** = *p* <0.0005; **** = *p* <0.00005. Dorsal is up in all images.

### Cell death is elevated in the *dnmt1*^-/-^ CMZ in a *p53*-independent manner

Previous publications have demonstrated increased *p53* expression and TUNEL^+^ cells in Dnmt1-deficient tissues and cell types^30,34,35,47^ suggesting a *p53*-dependent apoptotic mechanism for cell loss. Based on these studies, we hypothesized that *dnmt1*^-/-^ RSCs would similarly undergo p53-dependent apoptosis. To test this hypothesis, we first assayed for the presence of DNA double-strand breaks in *dnmt1*^-/-^ and sibling retinae using TUNEL (Figure 2A-F). *dnmt1* siblings displayed few TUNEL^+^ cells between 3-5dpf (Figure 2L-N), whereas the *dnmt1*^-/-^ retina contained increased proportions of TUNEL^+^ cells at 3, 4, and 5dpf in the INL (+0.5-2.3%, *p* <0.05), ONL (+0.01-1.8%, *p* <0.05) and at 5dpf in the GCL (+1.3%, *p* <0.05; Figure 2H-I). Within the CMZ, we detected a 4.5% increase in TUNEL^+^ cells at 3dpf (*p* <0.005, Figure 2J) prior to the onset of CMZ disorganization. This proportion decreased to 1% at 4dpf (*p* <0.05) and increased again to 3.7% at 5dpf (*p* <0.05; Figure 2J), a time at which *dnmt1*^-/-^ larvae begin to display severe systemic defects. During this 3-5dpf period, the majority of TUNEL^+^ cells in *dnmt1*^-/-^ larvae were located within the retina proper, not within the CMZ (Figure 2K and Supplemental Fig S2). In concordance with the TUNEL data, immunofluorescence of the pro-apoptotic marker, active-caspase3, displayed similar patterns to TUNEL (data not shown). Together, these data are consistent with those seen in previous studies; dnmt1 deficiency results in increased cell death^30,35,47^.

**Figure 2.**
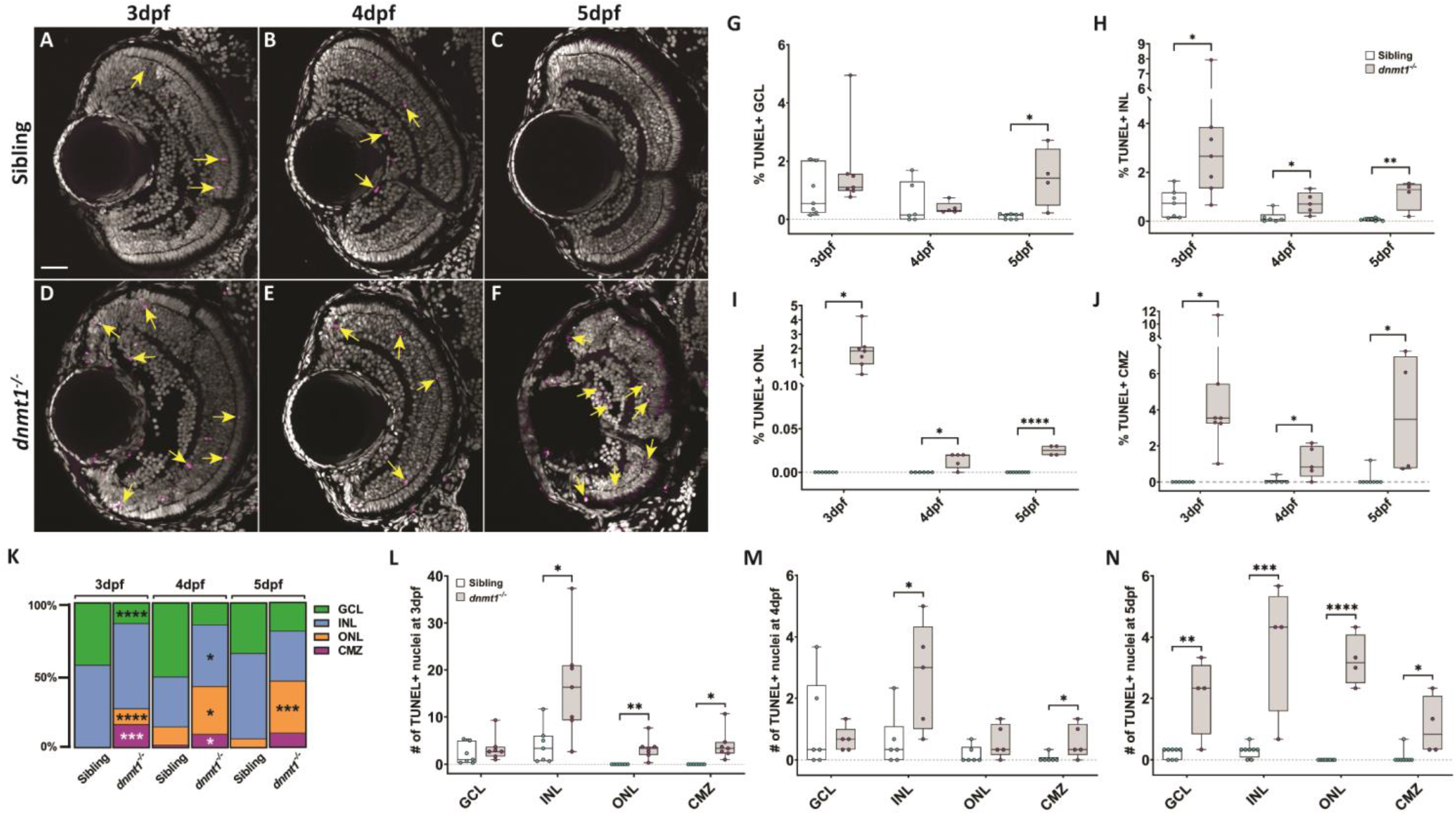
Cell death is elevated in the *dnmt1*^-/-^ CMZ. A-F. dnmt1 sibling (**A-C**) and mutant (**D-F**) retinae labeled with DAPI
(gray; nuclei) and TUNEL (magenta; dsDNA breaks) from 3-5dpf. **G-J.** Proportion of retinal layers (GCL, INL, ONL, and CMZ) labeled by TUNEL staining. **K.** Proportion of TUNEL^+^ cells within each layer from 3-5dpf. **L-N.** Average number of TUNEL^+^ cells in each retinal layer of siblings and *dnmt1*^-/-^ larvae from 3-5dpf. Yellow arrows in **A-F** indicate TUNEL^+^ nuclei. Scale bars = 30 μm. \**p* <0.05, \*\**p* <0.005, \*\*\**p* <0.0005, \*\*\*\**p* <0.00005. Dorsal is up in all images.

To identify if dnmt1 deficient RSCs are lost via *p53*-dependent apoptosis, we generated *dnmt1;p53* double mutants using the *p53^zdf1^* allele, which is defective in *p53*-dependent apoptosis^48,49^. We hypothesized that *p53*-dependent apoptosis was the driving mechanism of RSC loss in *dnmt1*^-/-^ mutants and therefore loss of p53 activity would rescue the CMZ phenotype. To test this hypothesis, we quantified nuclei in *dnmt1*^+/+^;*p53*^+/+^, *dnmt1*^+/+^;*p53*^-/-^, *dnmt1*^-/-^;*p53*^+/+^ and *dnmt1*^-/-^;*p53*^-/-^ retinae (Figure 3). Loss of p53 function did not affect retinal morphology (Figure 3A,E,I compared to Figure 3B,F,J) and *dnmt1*^+/+^; *p53*^-/-^ mutants possessed equivalent retinal cell numbers as *dnmt1*^+/+^;*p53*^+/+^ siblings (Figure 3Q-T and Supplemental Fig S2D) at 3,4 and 5dpf. When considering *dnmt1*^-/-^;*p53*^-/-^ larvae, we predicted an increase in CMZ cell numbers and a rescue of the CMZ-specific phenotype when compared to *dnmt1*^-/-^; *p53*^+/+^ larvae. Surprisingly, the *dnmt1^-/-^*;*p53*^-/-^ CMZ displayed similar morphology (Figure 3C,D,G,H,K,L) and was proportional to the *dnmt1*^-/-^; *p53*^+/+^ sibling retina (Figure 3P) across all three time points. These results suggest that *p53*-dependent apoptosis is not responsible for *dnmt1^-/-^* RSC loss.

**Figure 3.**
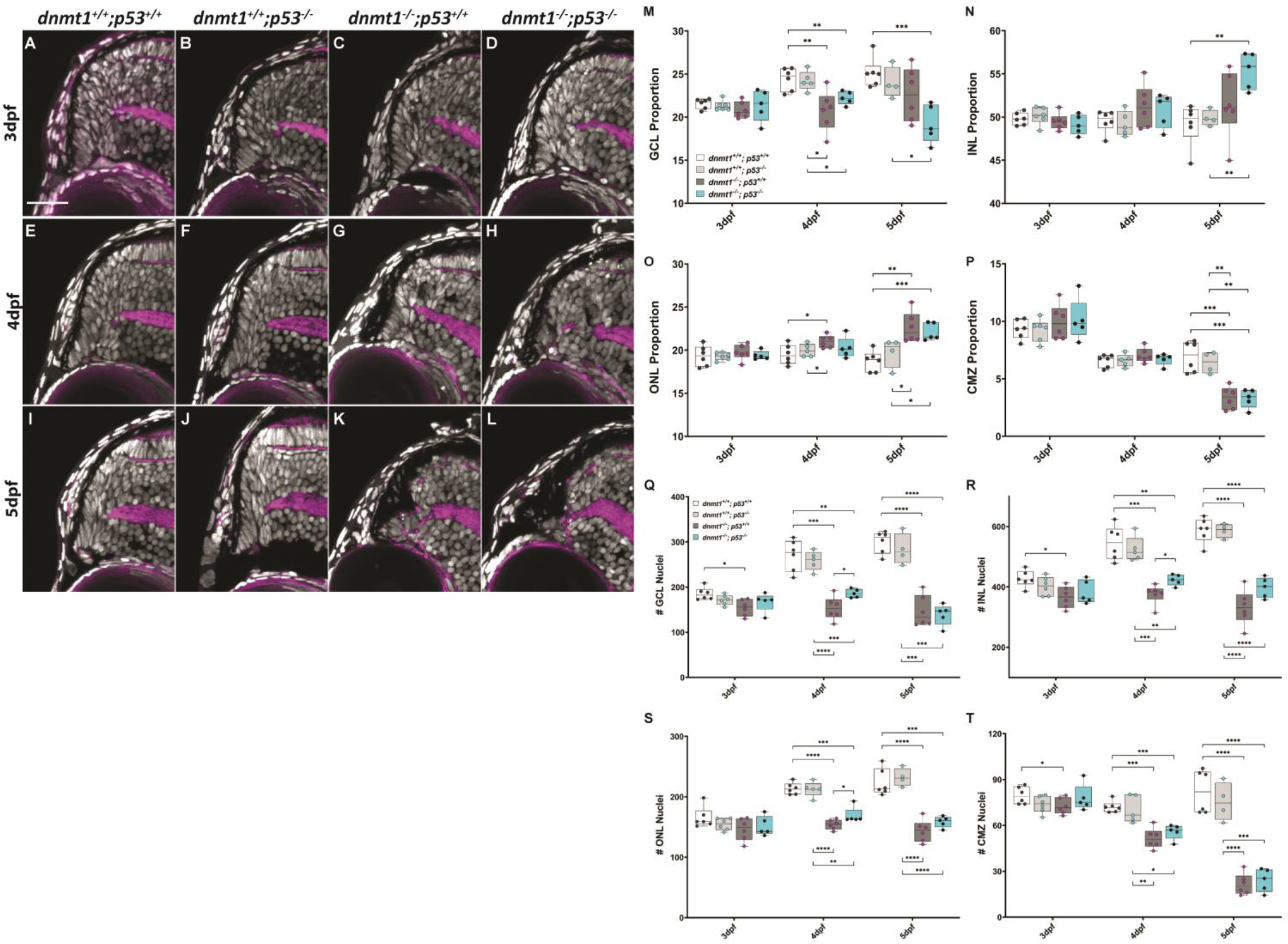
Loss of p53 function does not rescue the *dnmt1*^-/-^ CMZ phenotype. **A-L.** Transverse sections of the dorsal CMZ in wildtype (**A, E, I**), *dnmt1*^+/+^;*p53*^-/-^ (**B, F, J**), *dnmt1^-/-^;p53^+/+^* (**C, G, K**), *dnmt1*^-/-^;*p53*^-/-^ (**D, H, L**) larvae from 3-5dpf. Nuclei labeled with DAPI (gray) and F- actin labeled with phalloidin (magenta). **M-P.** Graphs depicting changes in retinal domain proportions over time. **Q-T.** Number of nuclei in each retinal domain of *dnmt1*^+/+^;*p53*^+/+^, *dnmt1*^+/+^;*p53*^-/^, *dnmt1*^-/-^;*p53*^+/+^, and *dnmt1*^-/-^;*p53*^-/-^ larvae from 3-5dpf. GCL: ganglion cell layer; INL: inner nuclear layer; ONL: outer nuclear layer; CMZ: ciliary marginal zone. Scale bars = 25 μm. \**p* <0.05; \*\**p* <0.005; \*\*\**p* <0.0005; \*\*\*\**p* <0.00005. Dorsal is up in all images.

### *dnmt1* is required to maintain RSC gene expression

*dnmt1* is expressed in RSCs at 4dpf (Figure 4I,J), consistent with dnmt1’s known requirements in stem cell populations *in vivo*^27,30,31,50,51^. Loss of Dnmt1 function results in aberrant gene expression in a number of contexts^45,50,52,53^ and therefore we wanted to determine if CMZ-specific gene expression was altered in the *dnmt1*^-/-^ CMZ. Previous reports have characterized the expression/distribution of the CMZ-specific genes: *col15a1b*, *cyclinD1*, *cdkn1c*, and *atoh7*^14,15,54^. To determine if CMZ expression of these genes was altered in *dnmt1*^-/-^ larvae, we utilized whole-mount *in situ* hybridization at 4dpf when the morphological defects in the CMZ begin to manifest (Figure 1). All sibling controls displayed normal CMZ-specific gene expression at 4dpf (Figure 4). Expression of *col15a1b* and *atoh7* were normal in *dnmt1*^-/-^ larvae (Figure 4C,D,S,T); however, the expression of *ccnD1* and *cdkn1ca*, which function to regulate cell cycle progression, were disrupted (Figure 4G,H,O,P). The majority of 4dpf *dnmt1*^-/-^ CMZs maintained *dnmt1* expression (Figure 4K,L). These data suggest that RSCs are present at the onset of morphological defects in the *dnmt1*^-/-^ CMZ, but could be impaired in their ability to progress through the cell cycle and self-renew.

**Figure 4.**
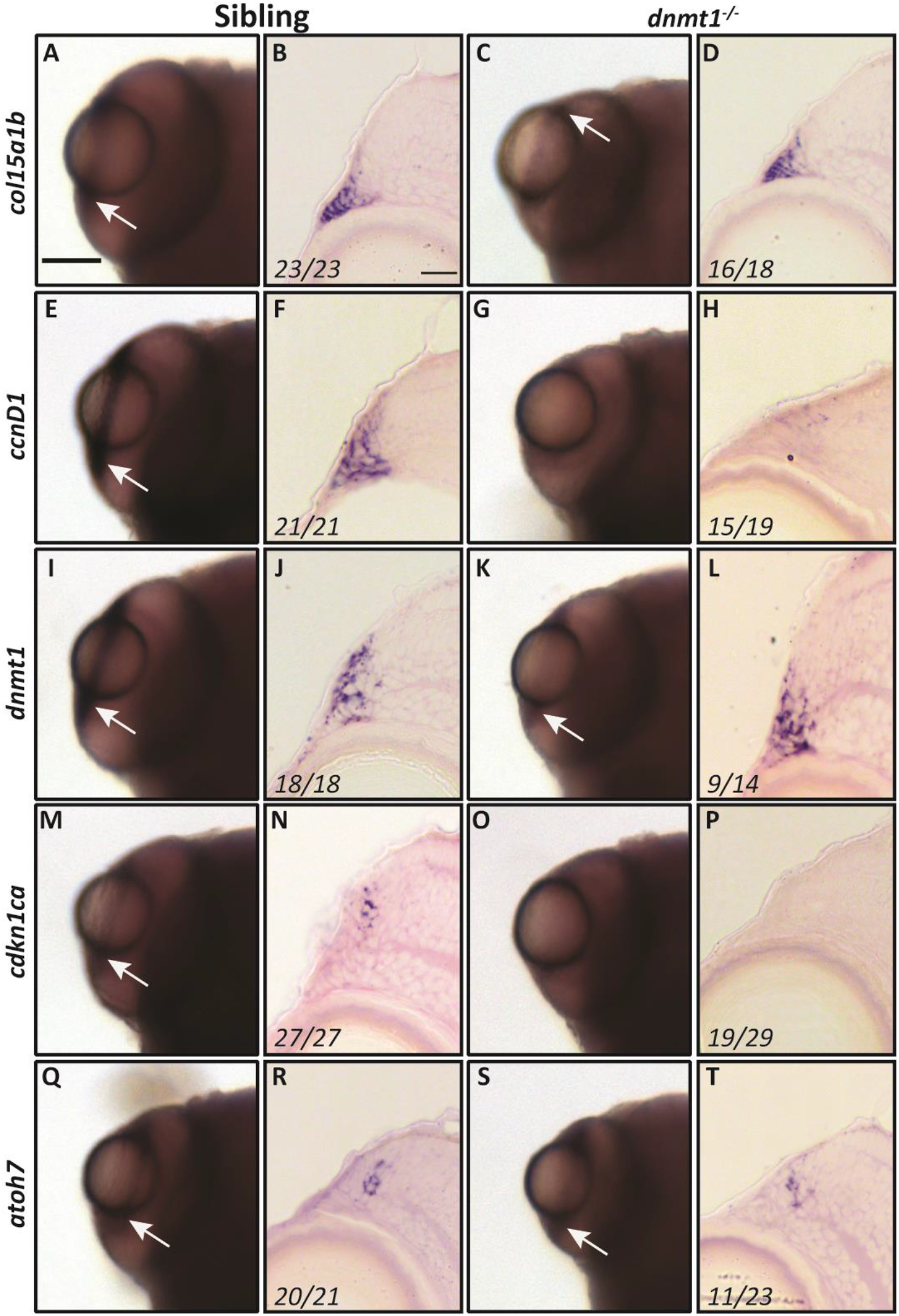
dnmt1 is required to maintain RSC gene expression. Gene expression shown in whole mount (**A, C, E, G, I, K, M, O, Q, S**) and transverse cryosections (**B, D, F, H, J, L, N, P, R, T**) between siblings and *dnmt1^-/-^ larvae*. **A-D.** *col15a1b* expression. **E-H.** *ccnD1* expression. **I-L.** *dnmt1* expression. **M-P.** *cdkn1ca* expression. **Q-T.** *atoh7* expression. Numbers in transverse cryosections designate the number of larvae that showed the displayed expression pattern vs. the total number of larvae analyzed. Scale bars = 75 mm (whole mount) and 10 μm (transverse sections). Anterior is up in all whole-mounts and dorsal is up for all section images.

### Loss of *dnmt1* activity results in decreased RSC proliferation

RSCs within the teleost CMZ remain proliferative throughout the lifespan of the animal^3,55,56^ and Dnmt1 is known to be required for cell cycle progression within stem cells of various tissue types^24,25,57^. Based on the significant loss of RSCs in *dnmt1*^-/-^ larvae between 3-5dpf (Figure 1) and the inability of *dnmt1*^-/-^ RSCs to maintain expression of cell cycle genes (Figure 4), we hypothesized that *dnmt1*^-/-^ RSCs would be defective in their proliferative capacity. To test this hypothesis, larvae were incubated for 2 hours in BrdU at 3, 4, and 5dpf, fixed immediately thereafter, and immunolabeled for BrdU and phosphohistone-H3-serine10 (pH3) to identify RSCs in late G2/M. *dnmt1* siblings maintained a constant proportion of BrdU^+^ cells within the CMZ between 3-5dpf (Figure 5A-C,H). Notably, the proportion of BrdU^+^ *dnmt1*^-/-^ RSCs at 3dpf was comparable to sibling controls (compare images in Figure 5A and 5D and nuclear proportions in Figure 5G). However, beginning at 4dpf, the percentage of BrdU^+^ *dnmt1*^-/-^ RSCs is significantly reduced when compared to controls (Figure 5B,E,H; *p* <0.00001), and this proportion continues to decrease through 5dpf (Figure 5C,F,H; *p* <0.0001). Additionally, the proportion of cells in late G2/M phase (pH3^+^) was significantly reduced at 3 and 4dpf in the *dnmt1*^-/-^ CMZ when compared to siblings (Figure 5G,I) indicating potential cell cycle defects in *dnmt1*^-/-^ RSCs that manifest as early as 3dpf.

**Figure 5.**
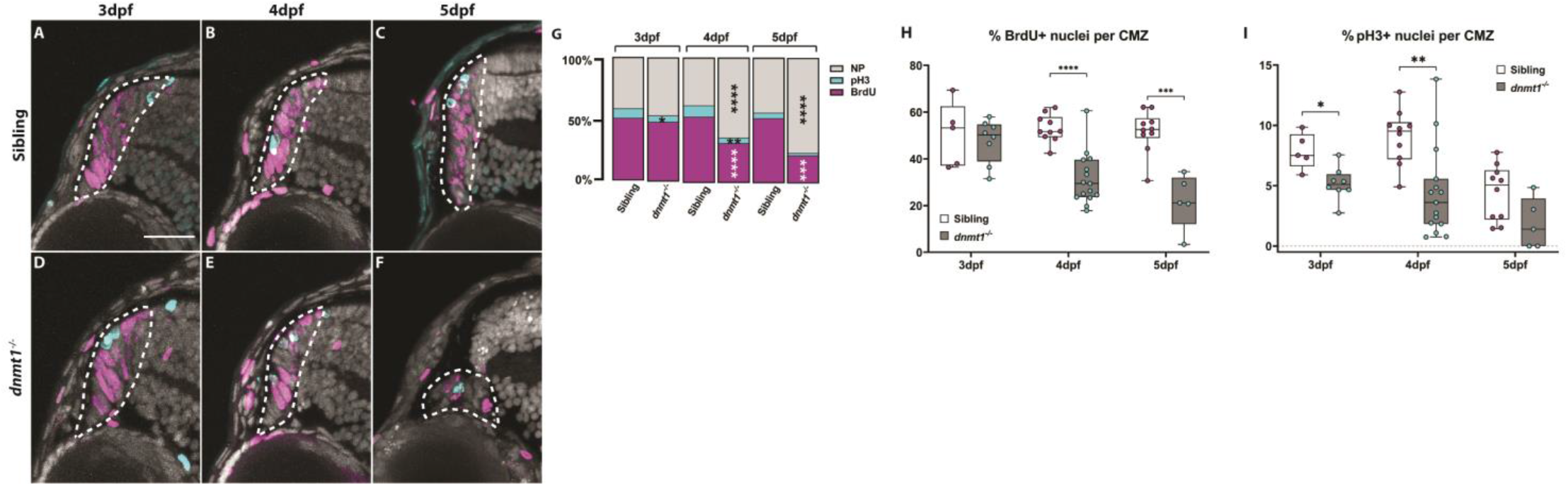
RSCs require dnmt1 function to maintain proliferation. A-F. Transverse sections of siblings (**A-C**) and *dnmt1*^-/-^ (**D-F**) larvae from 3-5dpf. Nuclei labeled with DAPI (gray). Cells in S-phase indicated by BrdU incorporation (magenta). Mitotic cells are labeled by a pH3(ser10) antibody (cyan). **G.** Proportions of CMZ cells in S-phase (magenta), G2/M-phase (cyan), or not proliferating (gray) of both siblings and *dnmt1*^-/-^larvae from 3-5dpf. **H.** Proportion of CMZ cells labeled with BrdU from 3-5dpf between controls and *dnmt1*^-/-^larvae. **I.** Proportion of CMZ cells labeled with pH3 from 3-5dpf between controls and *dnmt1*^-/-^larvae. White dotted lines designate CMZ (**A-F**). Scale bars: 30 μm \**p* <0.05, \*\**p* <0.005, \*\*\**p* <0.0005, \*\*\*\**p* <0.00005. Dorsal is up in all images.

### *dnmt1* is required for RSC differentiation and incorporation into the neural retina

Potential cell cycle progression defects coupled to the fact that the vast majority of *dnmt1*^-/-^ RSCs elude *p53*-dependent apoptosis (Figures 2 & 3) led us to hypothesize that *dnmt1*^-/-^ RSCs might instead be undergoing premature differentiation, as has been shown *in vitro*^28^. To test this hypothesis, we performed a BrdU birth-dating assay^58^. Our aim was to saturate RSCs with BrdU for a 12-hour period (3-3.5dpf) and quantify the average starting number of proliferating cells at 3.5dpf and determine the final position of daughter cells at 5dpf, once they incorporated into the retina (Figure 6A). Initial analysis of these samples revealed that most BrdU^+^ nuclei in both sibling and *dnmt1*^-/-^ larvae were located within the CMZ after the 12hr incubation (Figure 6C,E,G). However, there were a few BrdU^+^ cells that had incorporated into the neural retina at this time (Figure 6G). By comparing the number of BrdU^+^ nuclei of each retinal domain (CMZ, GCL, INL, ONL, Figure 6B) to the total number of BrdU^+^ nuclei (Figure 6H) at 3.5dpf, we noted a significant increase in the proportion of BrdU^+^ nuclei in the *dnmt1*^-/-^ CMZ (79.5%, *p* <0.05) compared to controls (71.7%, Figure 6G; Supplemental Fig. S3A). Additionally, we found that the proportion of BrdU^+^ cells in the *dnmt1*^-/-^ ONL (7.8%, *p* <0.05) was significantly reduced compared to siblings (10.9%, Figure 6G; Supplemental Fig. 3A) at 3.5dpf.

**Figure 6.**
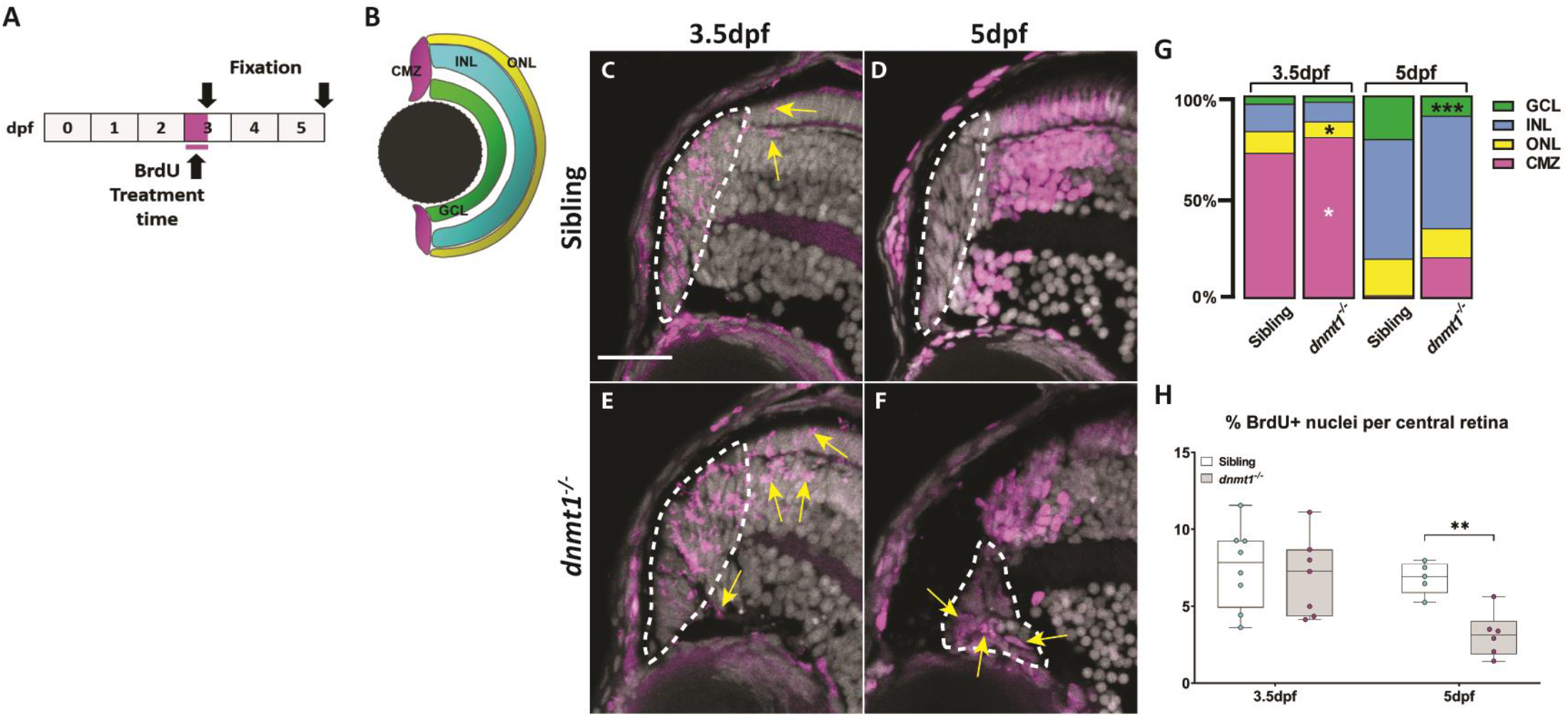
Neurons produced by *dnmt1*^-/-^ RSCs fail to integrate into the neural retina. **A.** Experimental paradigm depicting BrdU incorporation from 3-3.5dpf. Fixations occurred at 3.5 & 5dpf. **B.** Diagram of the four retinal domains (CMZ, GCL, INL, and ONL) whose colors correlate with the data presented in G. **C-F.** Transverse sections of BrdU pulses from 3-3.5dpf (C,E) and pulse-chase assay from 3-5dpf (D,F) (Siblings: A-B; *dnmt1*^-/-^ C-D). Nuclei labeled with DAPI (gray). Cells in S-phase indicated by BrdU incorporation (magenta). Mitotic cells are labeled by a pH3(ser10) antibody (cyan). **G.** Proportion of BrdU+ cells located in each retinal layer at 3.5dpf and 5dpf of *dnmt1*^-/-^ and control larvae. **H.** Proportion of total BrdU^+^ cells within the central retina of the pulse-chase experiment. White dotted lines designate the CMZ (**C-F**). Yellow arrows = BrdU^+^ nuclei outside the CMZ (**C,E**). Scale bars: 20 μm. \**p* <0.05, \*\**p* <0.005, \*\*\**p* <0.0005. Dorsal is up in all images.

At 5dpf, all BrdU^+^ cells in the sibling controls had exited the cell cycle and incorporated into the neural retina (Figure 6D,G), whereas *dnmt1*^-/-^ larvae retained 19.8% (*p* =0.05) of BrdU^+^ nuclei within the CMZ and had fewer BrdU^+^ cells overall within the retina (Figure 6F,G). Additionally, there was a significant decrease in the proportion of BrdU^+^ nuclei in the GCL (9.8%, *p* <0.0005) (Figure 6G; Supplemental Fig S3B) compared to controls (21.4%). Surprisingly, among the cells that remained in the 5dpf *dnmt1*^-/-^ CMZ, there was an increase in the BrdU^+^ proportion when compared to siblings (19.76% vs. 0.9% respectively, *p* =0.05; Figure 6G; Supplemental Fig S3B) suggesting an inability for some RSCs to either successfully complete the cell cycle or to integrate into retinal laminae. These data also show that daughter cells produced from the *dnmt1*^-/-^ CMZ proportionally incorporate into the INL and ONL at similar levels to those detected in controls (Figure 6G; Supplemental Fig S3B) supporting the notion that *dnmt1*^-/-^ RSCs are still capable of producing neurons that can successfully integrate into these two layers of the retina.

### Loss of dnmt1 activity leads to altered Long Terminal Repeat retroelement expression within the CMZ

Half of the zebrafish genome is comprised of endogenous viral elements known as transposons^59,60^, and dnmt1 is required for repressing the retroelement (RE) lineage of transposons^37,61–63^. Though many REs have lost their ability to “jump” throughout evolution, some still retain this ability^64,65^. These studies led us to hypothesize that aberrant DNA methylation resulting from loss of dnmt1 activity in RSCs would result in upregulation of RE expression within the *dnmt1*^-/-^ CMZ. To identify RE expression within the CMZ, we performed *in situ* hybridizations targeting several REs that belong to the Long Terminal Repeat (LTR) class of retrotransposons, specifically *Bel20*, *ERV1*, *ERV1-N5*, *ERV4*, and *Gypsy10* LTRs. We noted endogenous expression of *Bel20*, *ERV4*, and *Gypsy10* REs within the CMZ but not the neural retina of control larvae at 4dpf (Figure 7A,D,E). This result was unexpected since REs can be deleterious to cellular function^37,66–68^. However, not all of the LTR REs were detected within control CMZs; *ERV1* and *ERV1-N5* expression was not detected in the CMZ of siblings (Figure 7B,C), but rather *ERV1-N5* seemed to be expressed within the ONL of some control larvae (Supplemental Fig. S4O). Remarkably, *dnmt1*^-/-^ larvae had increased expression of *ERV1-N5* in the CMZ and within the overlying retinal pigmented epithelium (Figure 7H). The expression domains of *Bel20* and *ERV4* were expanded beyond the CMZ and into the neural retina of *dnmt1*^-/-^ larvae (Figure 7F,I) when compared to controls. Of note, we also identified several non-ocular tissues that displayed altered RE expression between *dnmt1*^-/-^ and sibling control larvae (Supplemental Fig S4). Interestingly, these LTR RE expression patterns were larvae-dependent, suggesting that not all RSCs respond uniformly to loss of dnmt1 function.

**Figure 7.**
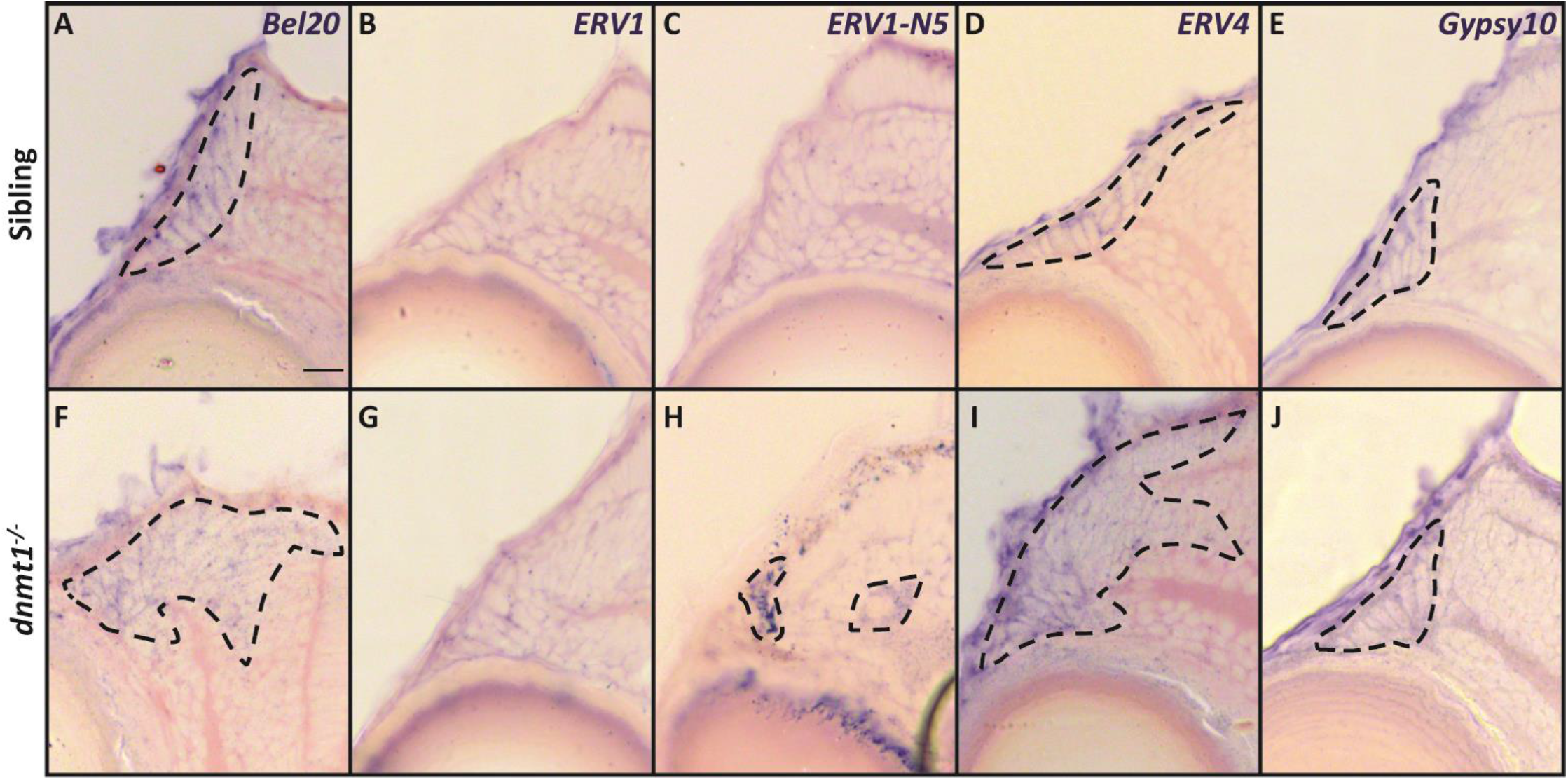
Loss of dnmt1 function results in misregulation of retroelement expression. **A-J.** Transverse cryosections of sibling (**A-E**) and *dnmt1*^-/-^ (**F-J**) larvae at 4dpf. **A, F.** Expression of *Bel20 LTR*. **B, G.** Expression of *ERV1 LTR*. **C, H.** Expression of *ERV1-N5 LTR*. **D, I.** Expression of *ERV4 LTR*. **E, J.** Expression of *Gypsy10 LTR*. Dotted lines: domains of retroelement expression. Scale bars = 10 μm. Dorsal is up in all images.

### A *L1RE3-EGFP* transgene reports increased *LINE1* retrotransposition activity in *dnmt1*^-/-^ CMZ

To expand our analysis of RE expression in *dnmt1*^-/-^ RSCs, and more specifically, visualize retrotransposition activity *in vivo*, we generated a non-LTR, *LINE1* element transgenic reporter line by modifying the *pLRE3-EGFP* plasmid^69,70^ (referred to as *L1RE3-EGFP* for the remainder of this study). The *L1RE3-EGFP* construct contains a human-derived *LINE1* RE sequence that requires retrotransposition for EGFP to be expressed and translated into a functional protein^69^. p53 is known to repress REs and when used transiently in *p53*^-/-^ zebrafish, *L1RE3-EGFP* was shown to have increased transposition activity and EGFP expression^68^. We validated the stability and effectiveness of the *L1RE3-EGFP* transgenic using again *p53* mutants^48,68^ and immunolabeling for EGFP (Supplemental Fig S5A). When *L1RE3-EGFP* was incorporated into the *dnmt1^s872^* genetic background, ectopic EGFP expression could be seen within the *dnmt1*^-/-^ eye when compared to control siblings (Supplemental Fig S5B,C). Notably, we detected ectopic EGFP expression more frequently within the *dnmt1*^-/-^ CMZ at both 3dpf (Figure 8C) and 4dpf (Figure 8D) timepoints when compared to controls (Figure 8A,B). However, like RE expression, clonal EGFP expression patterns were variable, both within and between sibling controls and *dnmt1*^-/-^ larvae, again suggesting that the effects of dnmt1 loss is variable from cell to cell and larva to larva.

**Figure 8.**
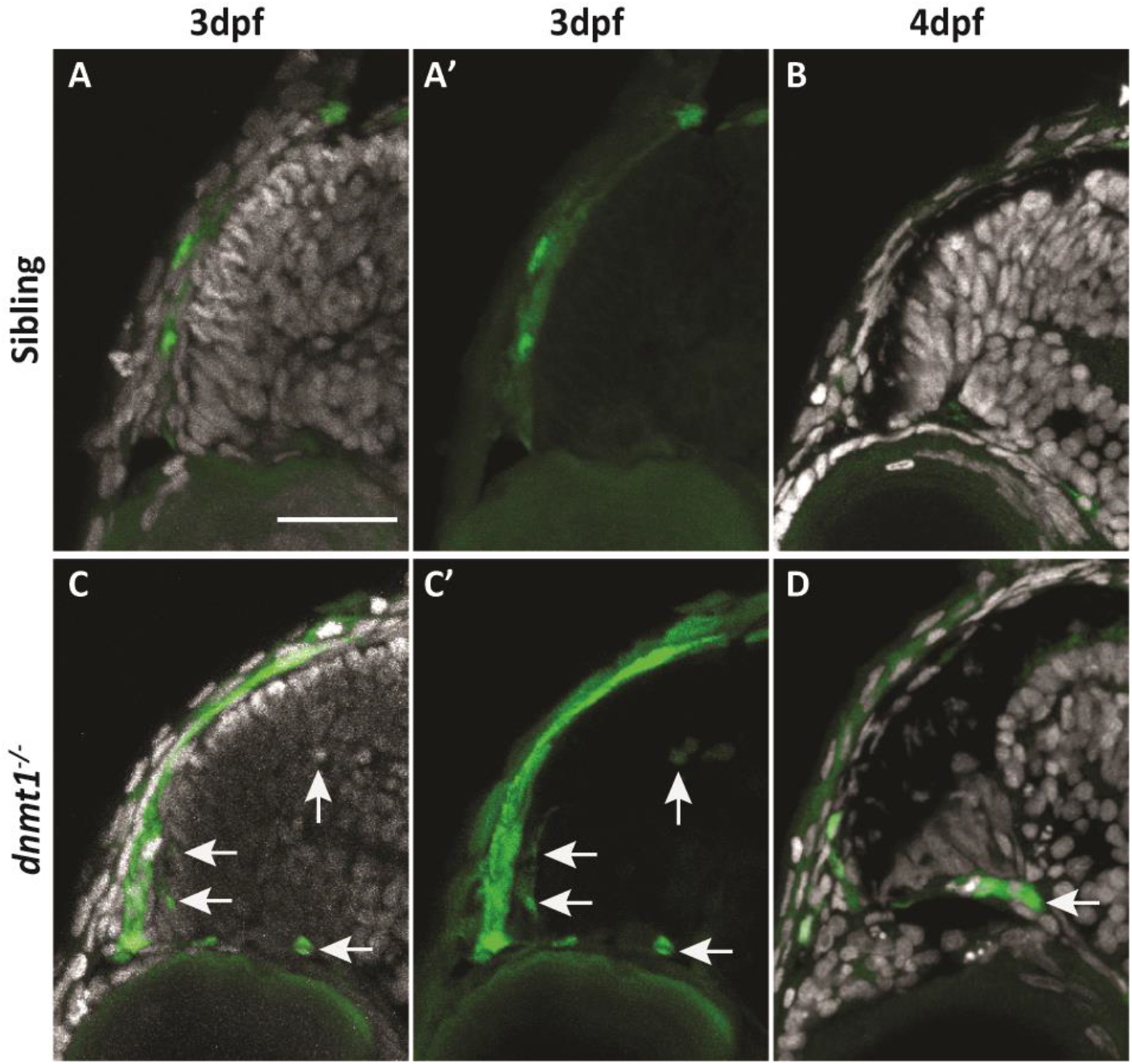
RSCs require dnmt1 function to repress L1RE3-EGFP transposition. **A-D.** Transverse sections of *Tg*(*CMV:Hsa.L1RE3,EGFP,myl7:EGFP;dnmt1*^+/+^) (**A-B**) and *Tg(CMV:Hsa.L1RE3,EGFP,myl7:EGFP;dnmt1^-/-^)* (**C-D**) larvae at 3dpf (**A, A’, C, C’**) and 4dpf (**B, D**). Nuclei labeled with DAPI (gray). Endogenous EGFP expression activated after L1RE3-EGFP transposition labeled in green. Arrows delineate EGFP+ cells. Scale bars: 30 μm. Dorsal is up in all images.

## Discussion

The zebrafish, with its lifelong, actively cycling RSCs within the CMZ, is a powerful model through which we can address how epigenetic regulators function to maintain these stem/progenitor cell populations *in vivo*. This study focused on the role of the DNA maintenance methyltransferase, *dnmt1*, within the CMZ, with the goal of determining how dnmt1 activity facilitates RSC maintenance. Previous work has shown that loss of dnmt1 function results in ocular defects^27,45,46,52^, but no studies have yet analyzed RSC populations and determined whether dnmt1 activity modulates their behavior.

Here, we demonstrate that *dnmt1* is essential for RSC homeostasis by maintaining CMZ-specific gene expression (Figure 4), facilitating cell cycle progression (Figure 5), and incorporation of CMZ-derived cells into the retina (Figure 6). These data are consistent with Dnmt1 functions described in other *in vivo* progenitor models such as the lens^27^, hippocampus^50^, kidney^62^, pancreas^30^ and intestine^51^. RSCs in S-and G2/M-phases of the cell cycle were detected in reduced proportions in the *dnmt1*^-/-^ CMZ and this correlated with a reduction in CMZ expression of genes encoding proteins that function in cell cycle progression, namely *ccnD1* (Figure 4G-H) and *cdkn1ca* (Figure 4O-P). Defects in cell cycle progression may also contribute to aberrant daughter cell integration into retinal laminae detected in *dnmt1*^-/-^ larvae (Figure 6). Although our data demonstrate an overall loss of RSCs and their progeny, this RSC depletion is independent of the p53-driven apoptotic pathway (Figures 2 and 3) signifying additional mechanisms of cell loss are at play.

While loss of *p53* function in the *dnmt1*^-/-^ background significantly rescued cell death within the laminated retina, validating that the *p53^zdf1^* allele is in fact inhibiting *p53*-driven apoptosis, loss of p53 in the *dnmt1*^-/-^ CMZ had no effect on CMZ cell numbers suggesting a *p53*-independent cell death pathway is likely modulated by dnmt1 in the CMZ^71^. Recent reports have demonstrated an upregulation of an innate inflammatory response in *dnmt1*^-/-^ larvae^37^. Necroptosis, a programmed cell death pathway tightly linked to a cell’s innate viral detection system and inflammatory responses, also results in DNA fragmentation and, in its later stages, is detected by TUNEL ^71^. Accordingly, we considered the possibility that *dnmt1*^-/-^ RSCs were instead lost via necroptosis. We tested this hypothesis using several chemical inhibitors of necroptosis, some of which have been reported to function in the zebrafish^72,73^; however, we were unable to replicate necroptotic inhibition nor validate drug efficacy. None the less, we predict that either necroptosis or pyroptosis (a programmed cell death pathway triggered by intracellular bacterial infections^74,75^) are the most likely mechanisms of cell death in dnmt1-deficient RSCs, but this will require the development of new tools to enable further analysis.

Alterations in RE expression activity the *dnmt1*^-/-^ CMZ (Figures 7,8) are exciting given Dnmt1’s known roles in repressing RE activity^36–39^. RE expression was aberrant in most *dnmt1*^-/-^ CMZs examined (Figure 7); however, expression changes and levels were variable between larvae, suggesting that the location and extent of genomic hypomethylation resulting from loss of dnmt1 function is inherently variable between cells of each larva. Previous reports demonstrated innate RE activity within somatic neural tissue^64,65,76–78^. Indeed, we detected retrotransposition activity within the larval zebrafish brain (Supplemental Fig S5D-I) of both siblings and *dnmt1*^-/-^ larvae from 2-4dpf, similar to activity detected in human hippocampal neurons^64,65,78^. However, RE retrotransposition is highly variable between larvae. Further studies will be required to determine what cellular processes might sensitize a cell-or tissue-type to upregulate REs and whether these REs have a mechanistic purpose within the cell.

In conclusion, our results demonstrate that dnmt1 functions to maintain RSC proliferation, gene expression, and integration of RSC daughters into the retina. Additionally, some REs are innately expressed within RSCs, however dnmt1 function is required to maintain tight control of these viral elements. Without dnmt1 activity, *LTR* expression remains active within the retina and *L1RE3-EGFP* retrotransposition activity is increased. Interestingly, RE activity within RSCs does not result in p53-mediated apoptosis, supporting a model in which *dnmt1*^-/-^ RSCs are lost through another mechanism of cell death. As discussed above, we predict that this increase in RE activity most likely activates necroptotic or pyroptotic cell death pathways, which are both known to result from intracellular responses to invading pathogens^71,74,75^. Regarding the innate *LTR* expression within *dnmt1*^+/+^ RSCs, in conjunction with previous reports of inherent RE activity within human neural tissue, it is worth considering how RE activity may contribute to neural stem cell biology. It is well known that dysregulation of REs is a hallmark of many human neurodegenerative diseases^67,79–82^. Future evaluations regarding the innate cost-to-benefit ratio of RE activity could provide crucial evidence for the development of neurodegenerative therapies.

## Methods

### Zebrafish maintenance

Zebrafish (*Danio rerio*) were maintained at 28.5°C on a 14 h light/ 10 h dark cycle. All protocols used within this study were approved by the Institutional Animal Care and Use Committee of The University of Pittsburgh School of Medicine, and conform to the National Institutes of Health Guide for the Care and Use of Laboratory Animals. Mutant alleles used in this study were *dnmt1^872^* and *tp53^zdf1^. dnmt1^s872^* and *tp53^zdf1^* zebrafish were genotyped using BioRad’s CFX Manager 3.1 and Precision Melt Analysis software (v4.0.52.0602). All genotyping primers are listed in Supplemental Table 1. Transgenic *Tg(CMV:Has.L1RE3,EGFP,myl7:EGFP)^pt701^* zebrafish were generated as described^83^ using constructs generously provided by Kristen Kwan and Chi-Bin Chien (University of Utah, Salt Lake City).

### BrdU labeling

To assess cellular proliferation, larvae were incubated in 10mM BrdU for either 2 or 12 hours, after which the BrdU was washed out and larvae were either collected or used for BrdU pulse-chase experiments.

### Immunohistochemistry and fluorescent labeling

Immunohistochemistry performed as described previously^84^. The following antibodies and dilutions were used: anti-BrdU antibody (Abcam, ab6326, 1:250), anti-phospho-histone H3 (Ser10) (EMD Millipore, 06-570, 1:250), anti-GFP (Thermo Fisher Scientific, A-11122, 1:50), goat anti-rat Cy3 secondary (Jackson Immuno Research, 112-165-003, 1:500), goat antirabbit Cy3 secondary (Jackson Immuno Research, 111-165-144, 1:500), and goat anti-rabbit Cy5 secondary (Jackson Immuno Research, 711-035-152, 1:500). Nuclei were counterstained with DAPI using Vectashield with DAPI (Vector Laboratories, H-1200). F-actin was labeled using AlexaFluor 633 Phalloidin (Thermo Fisher Scientific, 1:33, A22284). TUNEL-labeling was accomplished using TMR-Red In situ Cell Death Detection Kit (Sigma Aldrich, 12156792910).

### Cloning and Probe Synthesis

CMZ-specific probes have been published previously^14,27^. Retroelement probes were generated using reverse transcription-polymerase chain reaction (RT-PCR) on Trizol-isolated RNA from 24hpf and 5dpf embryos. Primer sequences were kindly provided by Dr. Kirsten Sadler (NYU Abu Dhabi) and PCR products were ligated into pGEM-T-easy vector (Promega Cat# PR-A1360) and verified by Sanger sequencing. Plasmids containing the correct clones were linearized and used as templates to *in vitro* transcribe digoxigenin-labeled RNA probes (Roche).

### In situ hybridization

Hybridizations using digoxigenin labeled antisense RNA probes were performed essentially as described^85^, except that they were pre-incubated with 1 mg/mL Collagenase type 1A (Sigma, C9891) to allow probe diffusion throughout the tissue. All probe primer sequences and plasmid construct information are listed in Supplemental Table 1.

### Microscopy and image processing

For sectioned embryos, imaging was performed with an Olympus FV1200 confocal microscope. Confocal Z-stacks were collected in 1μm optical sections. Z-stacks were max-projected using ImageJ (version 1.52r) software (National Institutes of Health) and quantification was conducted using the “Cell Counter” plugin. Figures were prepared using Adobe Illustrator CS6 (Adobe Systems). *In situ* cryosections were imaged utilizing a Leica DM2500 with a 100X oil immersion objective (NA: 1.25).

### Cell counting and quantification

Each data point was collected from an individual larva. Each larva was analyzed using three consecutive 12 μm sections of the central retina using the optic nerve and lens morphology as retinal landmarks. The average of the three consecutive sections was used as a single data point (n >4 for all datasets). Proportions of retinal domains were calculated by dividing the number of DAPI-labeled nuclei in each domain over the total number of retinal nuclei.

### Statistics

For all statistical analysis, data were imported into GraphPad Prism 8 software. Quantification of nuclei and immunolabeled cells was statistically assessed using Student’s two-tailed unpaired T-test with *p* <0.05 as a significance threshold.

### Generation of Tg(hLINE1-EGFP)

pLRE-mEGFPI plasmid was generously donated by Dr. John V. Moran (The University of Michigan School of Medicine)^69^. The Hsa.L1RE3-EGFP sequence was isolated from the pCEP4 backbone using NotI and SalI restriction enzymes and then inserted into pME-MCS plasmid from the Tol2 Gateway Kit. LR Clonase II Plus was used to carry out all Multisite Gateway assembly reactions^83^ using p5E-MCS (19ng), pME-Hsa.L1RE3-EGFP (77ng), p3E-polyA (19ng), and pDestTol2CG2 (103ng) plasmids. Capped Tol2 mRNA was synthesized from pCS2FA-transposase using the Ambion mMessage mMachine Sp6 in vitro transcription kit (Thermo Fisher Scientific, AM1340). Tol2 mRNA (75pg) was co-injected with pDEST-Hsa.L1RE3-EGFP (40 pg) into *dnmt1*^+/-^;*p53*^+/-^ incross embryos at the 1-cell stage. Embryos displaying acceptable levels of mosaic myl7:EGFP expression were raised to adulthood, and outcrossed to screen for founders. F1 embryos displaying ubiquitous myl7:EGFP expression were isolated and reared to generate the stable line *Tg(CMV:Has.L1RE3,EGFP,myl7:EGFP)^pt701^*.

## Acknowledgements

We thank members of the Gross lab, Elizabeth Phair, Tom Holkenborg and the University of Pittsburgh zebrafish community for helpful comments and suggestions on this work, and Dr. Hugh Hammer and the University of Pittsburgh Department for Laboratory Animal Research for fish maintenance. Dr. Stephen W. Wilson kindly provided the *col15a1b* plasmid. Dr. Kirsten Sadler kindly provided retroelement primer sequences for the *in situ* probes used in this study. Dr. John V. Moran kindly provided the *pLRE-mEGFPI* plasmid. We acknowledge support from NIH Grant RO1 EY29031 and NIH CORE Grant P30-EY08098 to the Department of Ophthalmology, the Eye and Ear Foundation of Pittsburgh, and from an unrestricted grant from Research to Prevent Blindness, New York, NY. Fish lines were obtained from the Zebrafish International Resource Center which is supported by the NIH.

## Contributions

J.M.G and K.M.A. designed and conceived the study; K.M.A. collected all samples and performed the experiments and analyses; J.M.G and K.M.A. interpreted the results, wrote and reviewed the manuscript.

