## Supplemental Information for "*dnmt1* function is required to maintain retinal stem cells within the ciliary marginal zone of the zebrafish eye"

*Supplementary Information:*

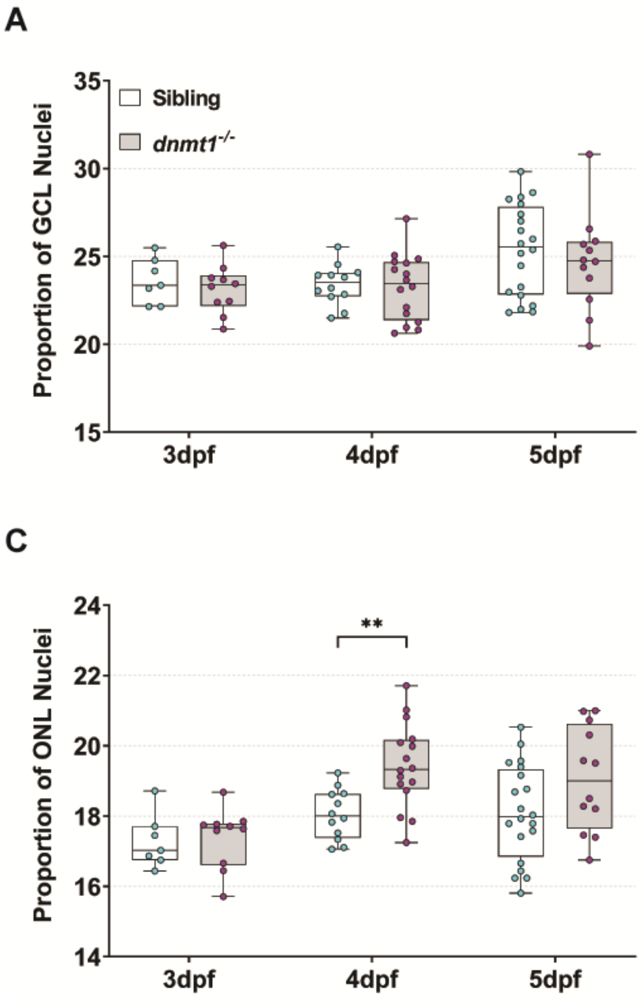


**Supplementary Figure S1. Loss of dnmt1 function results in a decline of RSC number. A-D.** Graphs of retinal domain proportions over time between siblings and *dnmt1*$^{-/-}$ larvae. **$p$ <0.005, *** = $p$ <0.0005.

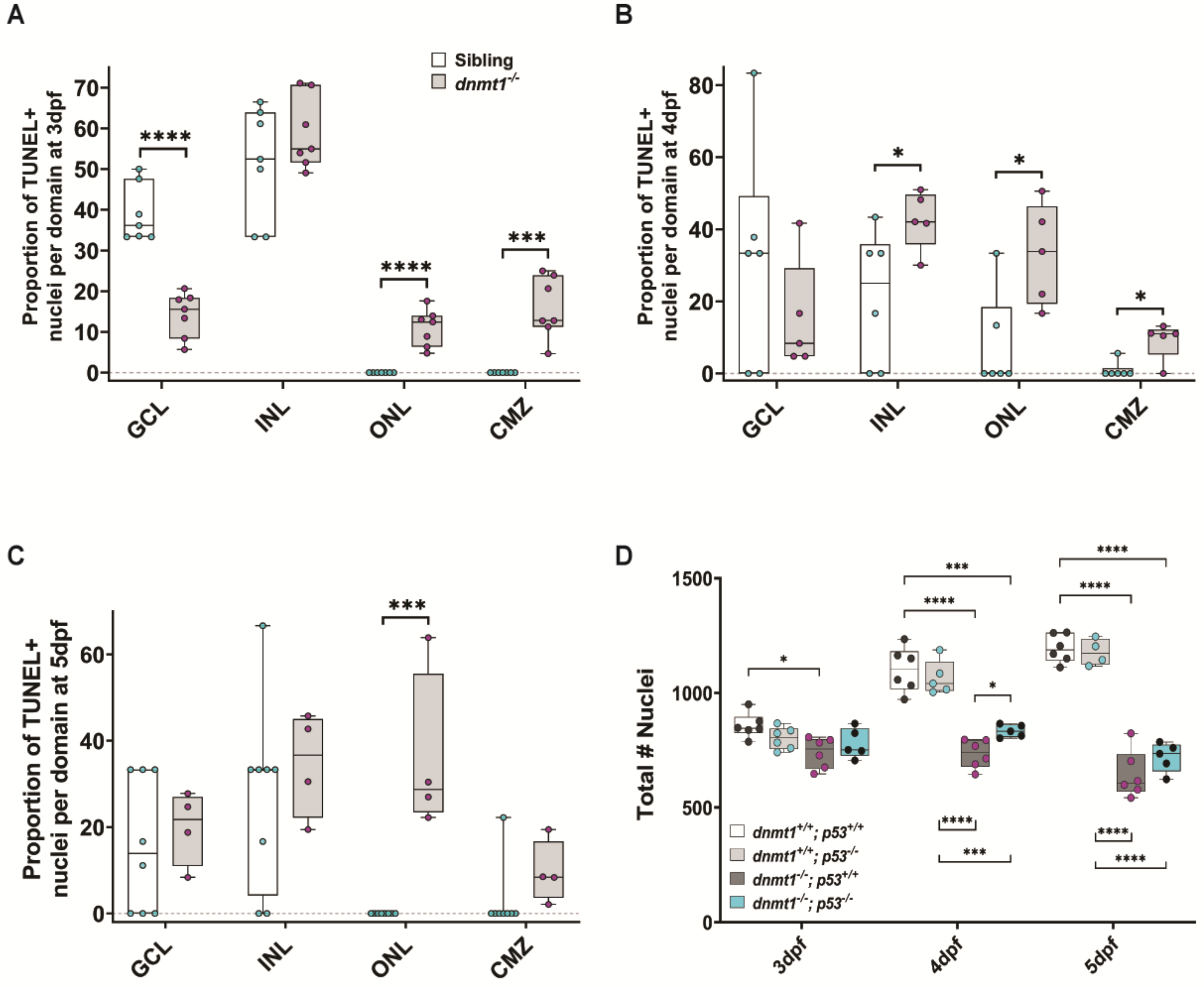


**Supplemental Figure S2. p53-mediated apoptosis is not responsible for *dnmt1*$^{-/-}$ RSC loss. A-C.** Proportion of TUNEL$^{+}$ nuclei in each retinal domain of *dnmt1*$^{-/-}$ larvae (gray bars) compared to sibling controls (white bars) at 3dpf (**A**), 4dpf (**B**), and 5dpf (**C**). D. Total number of retinal nuclei between *dnmt1*$^{+/+}$;*p53*$^{+/+}$ (white bars), *dnmt1*$^{+/+}$;*p53*$^{-/-}$ (light gray bars), *dnmt1*$^{-/-}$;*p53*$^{+/+}$ (dark gray bars), *dnmt1*$^{-/-}$;*p53*$^{-/-}$ (blue bars) larvae from 3-5dpf. GCL: ganglion cell layer; INL: inner nuclear layer; ONL: outer nuclear layer; CMZ: ciliary marginal zone. *$p < 0.05$, ***$p < 0.0005$, ****$p < 0.00005$. Dorsal is up in all images.

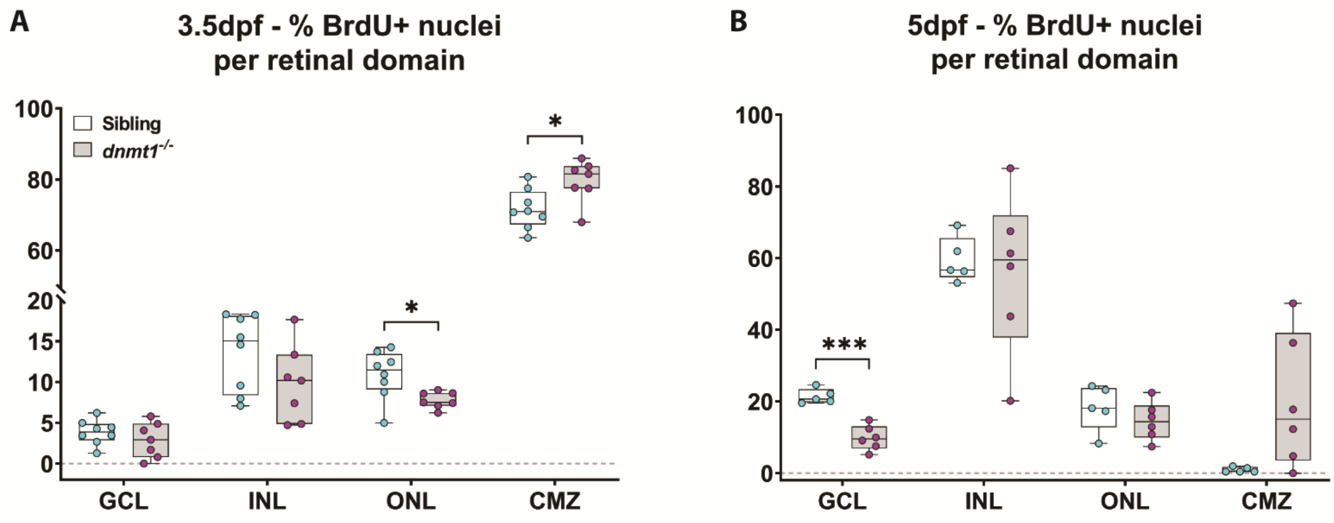


**Supplemental Figure S3. dnmt1-deficient RSCs fail to incorporate into the neural retina. A.** Data points collected of the proportion of cells labeled with BrdU in each retinal domain at 3.5dpf divided by the number of total BrdU$^+$ cells. **B.** Data points collected of the proportion of cells labeled with BrdU in each layer at 5dpf divided by the number of total BrdU$^+$ cells. Sibling controls = white bars; *dnmt1$^{-/-}$* = gray bars. Dorsal is up in all images.

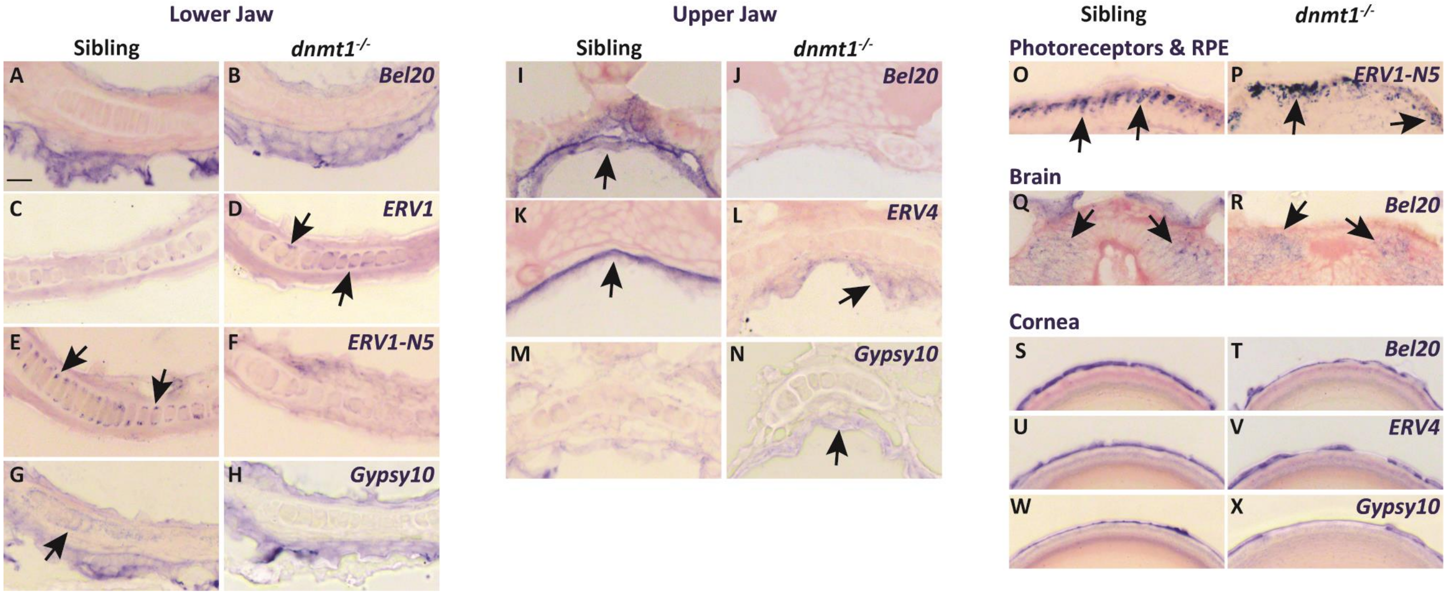


**Supplemental Figure S4. Loss of dnmt1 function results in misregulation of LTR RE expression across numerous tissues. A-X.** Transverse cryosections of larvae analyzed by *in situ* hybridization. All sibling and *dnmt1*$^{-/-}$ larvae are 4dpf. **A-H.** Expression of indicated REs within the lower jaw. **I-N.** Expression of indicated REs within the upper jaw. **O-P.** Expression of *ERV1-N5 LTR* in photoreceptors (**O**; sibling) and the RPE (**P**; *dnmt1*$^{-/-}$). **Q-R.** Expression of *Bel20 LTR* within the brain. **S-X.** Expression of indicated REs within the cornea. Arrows delineate expression changes of specified REs. Scale bar (A) = 10 µm. All images were taken at the same magnification. RPE = retinal pigmented epithelium.

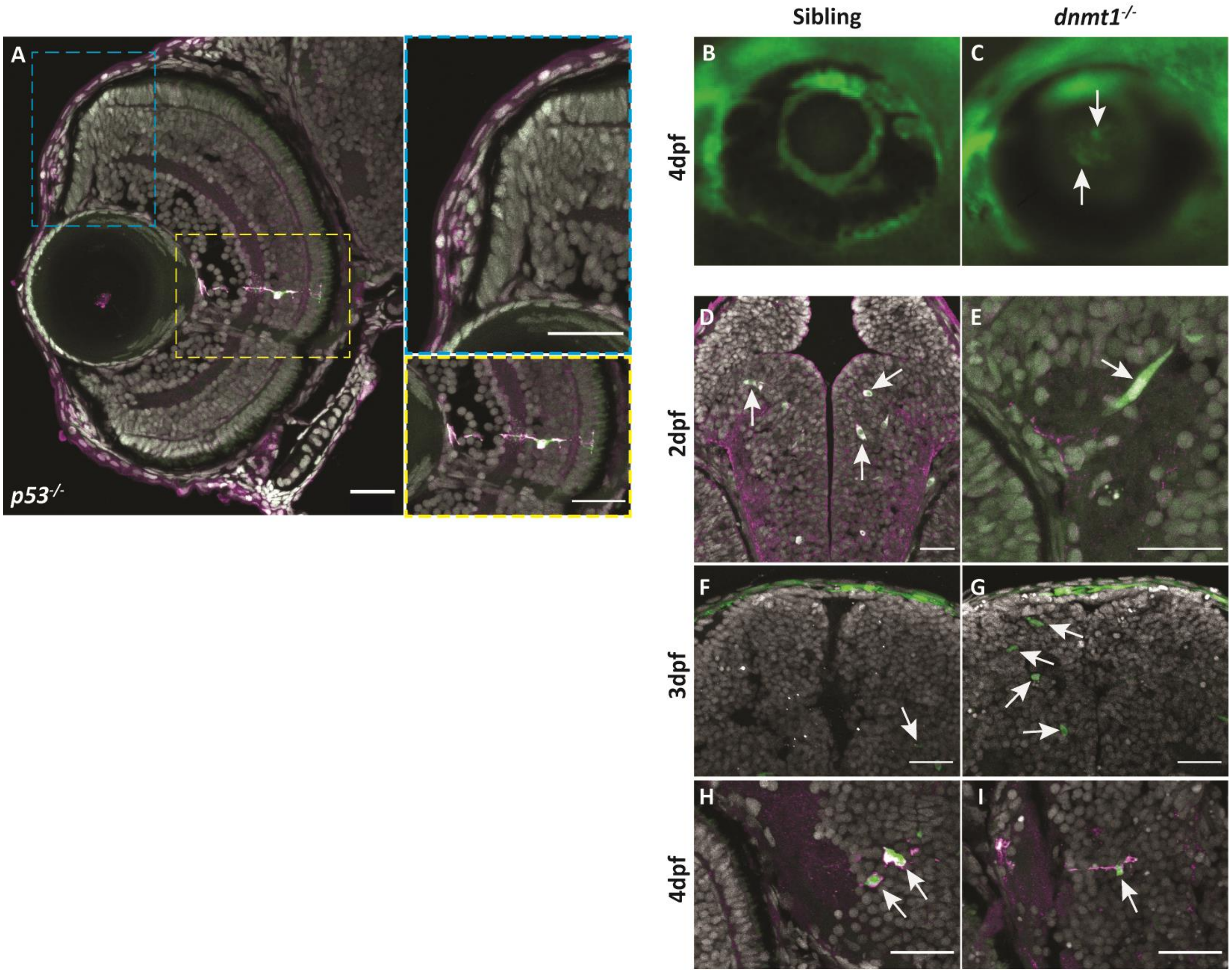


**Supplemental Figure S5. L1RE3-EGFP transgene expression is more prominent in *dnmt1$^{-/-}$* larvae. A-A''.** Transverse section of *Tg(CMV:Hsa.L1RE3,EGFP,myl7:EGFP;p53$^{-/-}$)* larvae at 4dpf. **A.** L1RE3-EGFP$^{+}$ retinal cells labeled with endogenous EGFP. Cyan (A') and yellow (A'') dotted boxes indicate magnified images to the right of panel A. **A'.** Magnified image of *Tg(CMV:Hsa.L1RE3,EGFP,myl7:EGFP;p53$^{-/-}$)* CMZ showing no expression of the L1RE3-EGFP transgene. **A''.** Magnified image of retinal neuron expressing L1RE3-EGFP transgene. **B-I.** All images are taken from *Tg(CMV:Hsa.L1RE3,EGFP,myl7:EGFP)* larvae that are either *dnmt1$^{+/+}$* (B,D,F,H) or *dnmt1$^{-/-}$* (C,E,G,I). **B-C.** Whole-mount images of 4dpf eyes demonstrating L1RE3-EGFP transgene activation seen through the lens of *dnmt1$^{-/-}$* larvae and not siblings. **D,F,H.** Sibling larvae expressing the L1RE3-EGFP transgene in the brain. **E,G,I.** *dnmt1$^{-/-}$* larvae expressing the L1RE3-EGFP transgene in the brain. Nuclei labeled with DAPI (gray). Endogenous L1RE3-activated EGFP labeled in green. EGFP antibody is magenta. Scale bars: 30 µm in all images. Dorsal is up in all images.

**Supplementary Table S1. List of primer sequences and appropriate experimental purpose used in this study.**

| ***Gene / Target*** | **Fwd Primer Sequence** | **Rev Primer Sequence** | **Experimental Purpose** |
|---|---|---|---|
| *dnmt1*$^{s872}$ | GAATCAATGCACACCATATGCC TTATGTAC | CTCAGGGTGAAGGACACGTCC | HRM Genotyping in TL strain |
| *dnmt1*$^{s872}$ | AATGCTGTTCCTCCACCCC | CTTGACCTCCAGGCCAATGG | HRM Genotyping in AB strain |
| *tp53*$^{zdf1}$ | CTAAACTACATGTGCAATAGCA GCTGC | CTCCAGAGTGATGATTGTGAGG ATGG | HRM Genotyping |
| *BEL20-LTR_DR* | AATGCAACGCAGTATCATCG | AGGTGCACTTCTCCGAGTGT | *In situ* hybridization probe; pGEM T-Easy Vector |
| *ERV1-1-LTR_DR* | TGAAGGATTTGATCTTCTCTCC | GCAATCGAGTCAGTGGGTCT | *In situ* hybridization probe; pGEM T-Easy Vector |
| *ERV1-N5-LTR_DR* | TGAGTATGCCTGCACAGGAG | TCAGACCGGAGGTTTTGAAT | *In situ* hybridization probe; pGEM T-Easy Vector |
| *ERV4_DR-I* | CCAAGACCGATCACACCTTT | ACTCCCATAATTCCCCCTTG | *In situ* hybridization probe; pGEM T-Easy Vector |
| *Gypsy10-LTR_DR* | TGCGGTTAACGCTTACAAAA | CACTCCCCCTAATCAGATACCA | *In situ* hybridization probe; pGEM T-Easy Vector |
